# Exploring protein sequence similarity with Protein Language UMAPs (PLUMAPs)

**DOI:** 10.1101/2022.09.27.509824

**Authors:** Adrian Jinich, Sakila Z. Nazia, Kyu Rhee

## Abstract

Visualizing relationships and similarities between proteins can reveal insightful biology. Current approaches to visualize and analyze proteins based on sequence homology, such as sequence similarity networks (SSNs), create representations of BLAST-based pairwise comparisons. These approaches could benefit from incorporating recent protein language models, which generate high-dimensional vector representations of protein sequences from self-supervised learning on hundreds of millions of proteins. Inspired by SSNs, we developed an interactive tool - Protein Language UMAPs (PLUMAPs) - to visualize protein similarity with protein language models, dimensionality reduction, and topic modeling. As a case study, we compare our tool to Sequence Similarity Network (SSN) using the proteomes of two related bacterial species, *Mycobacterium tuberculosis* and *Mycobacterium smegmatis*. Both SSNs and PLUMAPs generate protein clusters corresponding to protein families and highlight enrichment or depletion across species. However, only in PLUMAPs does the layout distance between proteins and protein clusters meaningfully reflect similarity. Thus in PLUMAPs, related protein families are displayed as nearby clusters, and larger-scale structures correlate with cellular localization. Finally, we adapt techniques from topic modeling to automatically annotate protein clusters, making them more easily interpretable and potentially insightful. We envision that as large protein language models permeate bioinformatics and interactive sequence analysis tools, PLUMAPs will become a useful visualization resource across a wide variety of biological disciplines. Anticipating this, we provide a prototype for an online, open source version of PLUMAPs.

## Introduction

With the drastic increase in accessibility to sequencing, there has been an explosion of available genomic data in the last two decades. The number of entries in the UniProtKB database continues to increase exponentially, with more than 12 million new entries alone since March 2021 ^1^. Despite the magnitude of new sequences identified, the rate of new annotations staggers behind the rate at which new sequences are identified. The great majority of these annotations are done computationally using sequence homology rather than manually through *in vivo* or *in vitro* experiments. Sequence homology groups proteins into superfamilies, clusters that share common sequence motifs, structural features, or biochemical substrates ^2^, guiding experimental exploration of unknown protein function.

The flood of protein sequence data calls for improvements in tools to visualize large protein collections. One popular computational approach to explore protein function through sequence homology is sequence similarity networks (SSN) ^3^. SSNs generate a measure of protein similarity by running BLAST-based pairwise alignments ^4^. Relationships between proteins are projected onto a multidimensional network, where each protein is a node connected to one or more proteins by a line if their sequence similarity exceeds a threshold specified by the user. By taking advantage of sequence similarity, SSNs allow visualization of much larger data sets and reveal relationships between more distant sequence sets in a protein superfamily that would have been obscured in multiple sequence alignment (MSA) or phylogenetic reconstructions. In conjunction with genomic and metagenomic information about proteins, SSNs are powerful tools in discovering enzyme functions ^5^.

While SSNs can provide significant information from BLAST-based sequence similarity, the recent field of protein language models promises to improve our ability to explore uncharacterized proteins. Protein language models harness machine learning methods from natural language processing (NLP) adapting them to proteins ^6^. The most powerful recent models use a transformer deep learning architecture, and are trained on large scale datasets (≈10^8^ proteins) with a self-supervised framework ^7–9^. In such self-supervised learning of large protein language models, the main training task consists of predicting randomly “masked” amino acids in sequences, giving the model a sense of residue interactions. As part of the learning algorithm, protein sequences, as well as individual amino-acids within each protein are encoded or embedded into vector representations of a fixed dimensionality.

Inspired by the practical value of Sequence Similarity Networks (SSN), here we present an open source and interactive protein visualization tool based on protein language models. Our pipeline uses the pre-trained ESM transformer language model ^7^ to generate vector representations for any set of input proteins. As similar proteins are encoded by the ESM close to each other in vector space, the protein representations are then clustered and projected to two dimensions. We then adapt techniques from topic modeling, a subfield of NLP that generates tools to uncover and visualize common underlying themes and narratives in language documents ^10–13^. We label each protein cluster with informative annotations by searching for enriched protein family terms and descriptors. To illustrate the value of our approach, named PLUMAPs, we visualize the proteome of the most important bacterial human pathogen, *Mycobacterium tuberculosis* and compare it to that of its non-pathogenic relative, *Mycobacterium smegmatis*. Similar to SSNs, we find that PLUMAPs tend to cluster proteins belonging to the same family together, therefore generating visually striking signals for protein enrichments or depletions between different proteomes. However, PLUMAPs preserve the global structure of protein similarity better than SSNs, with the larger-scale distances between proteins and protein clusters also containing meaningful information. This general notion is reflected in two specific findings: PLUMAPs group proteins by cellular localization (i.e. cytoplasmic vs. cell wall); some related protein family clusters are near each other in the visualization. Finally, additional layers of information can be overlaid on the PLUMAP, and we exemplify this by interactively displaying the AlphaFold2 structure of members of a protein cluster. We argue that PLUMAPs are an efficient visual analysis of protein sets, and are complementary to SSNs by helping non-trivial relationships between proteins come into focus [and generate insights.

## Results

### A protein language UMAP (PLUMAP) comparing the proteomes of *Mycobacterium tuberculosis* and *Mycobacterium smegmatis*

To illustrate the practical value of PLUMAPs, we focus as a case study on a visual comparison of two bacterial proteomes from our corner of microbiology. The slow growing *Mycobacterium tuberculosis* (Mtb) is the etiologic agent of tuberculosis (TB), the leading cause of death by bacterial infection. The fast growing and non-pathogenic *Mycobacterium smegmatis* (Msm, recently renamed as *Mycolicibacterium smegmatis*) is commonly used as a surrogate for Mtb in the laboratory. *Mtb* has a smaller proteome (≈4000 proteins) relative to Msm (≈6,600 proteins). While their two genomes have been extensively compared, PLUMAPs provide a fast and interactive visual representation that helps bring into focus insightful differences.

We generated a PLUMAP visualization of the *Mtb* and *Msm* proteomes via the following steps (Fig. 1) (see Methods for details). Using the pre-trained Evolutionary Scale protein language model (ESM-1b), which has a transformer architecture ^7^, we generated 1280-dimensional embeddings for all of *Mtb’s* and *Msm’s* proteins. We then clustered the protein vector representations and projected them to 2-dimensions with the UMAP: Uniform Manifold Approximation and Projection (UMAP) ^14^. Finally, using techniques from topic modeling [REF] and a natural language processing Python library [REF], we annotated the protein clusters with enriched annotation terms from the UniProt database. Finally, these annotated, low-dimensional projections of the protein language model embeddings can be interactively explored. Fig. 2A shows the composite PLUMAP for the *Mtb* and *Msm* proteomes. The general structure of the *Mtb/Msm* PLUMAP consists of two large protein islands and approximately 50-60 clearly distinct, smaller clusters. Interactive inspection of the smaller and clearly separated clusters shows that, like SSNs, the PLUMAP displays clusters that are enriched or depleted for Mtb or Msm proteins (Fig. 2B). As examples of known family enrichments, the map shows clusters of PE, PPE, and PE-PGRS families, as well as toxin-antitoxin families, which are enriched for *Mtb* proteins. *M. smegmatis* proteins are enriched in clusters of different families of redox enzymes, such as aldehyde dehydrogenases, a short-chain dehydrogenases, luciferase-like F420-dependent oxidoreductases, acyl-CoA dehydrogenases, and alcohol dehydrogenase-like proteins. Many clusters of transcriptional regulator families are enriched for *Msm* proteins, including TetR, MarR, and LysR families. A cluster of transporters of the Major Facilitator Superfamily (MFS) is visibly enriched for *Msm* proteins. These visually striking differential abundances can be used as preliminary guides for more in-depth protein family distribution profiles across larger sets of species (Fig. 2C).

**Fig. 1:**
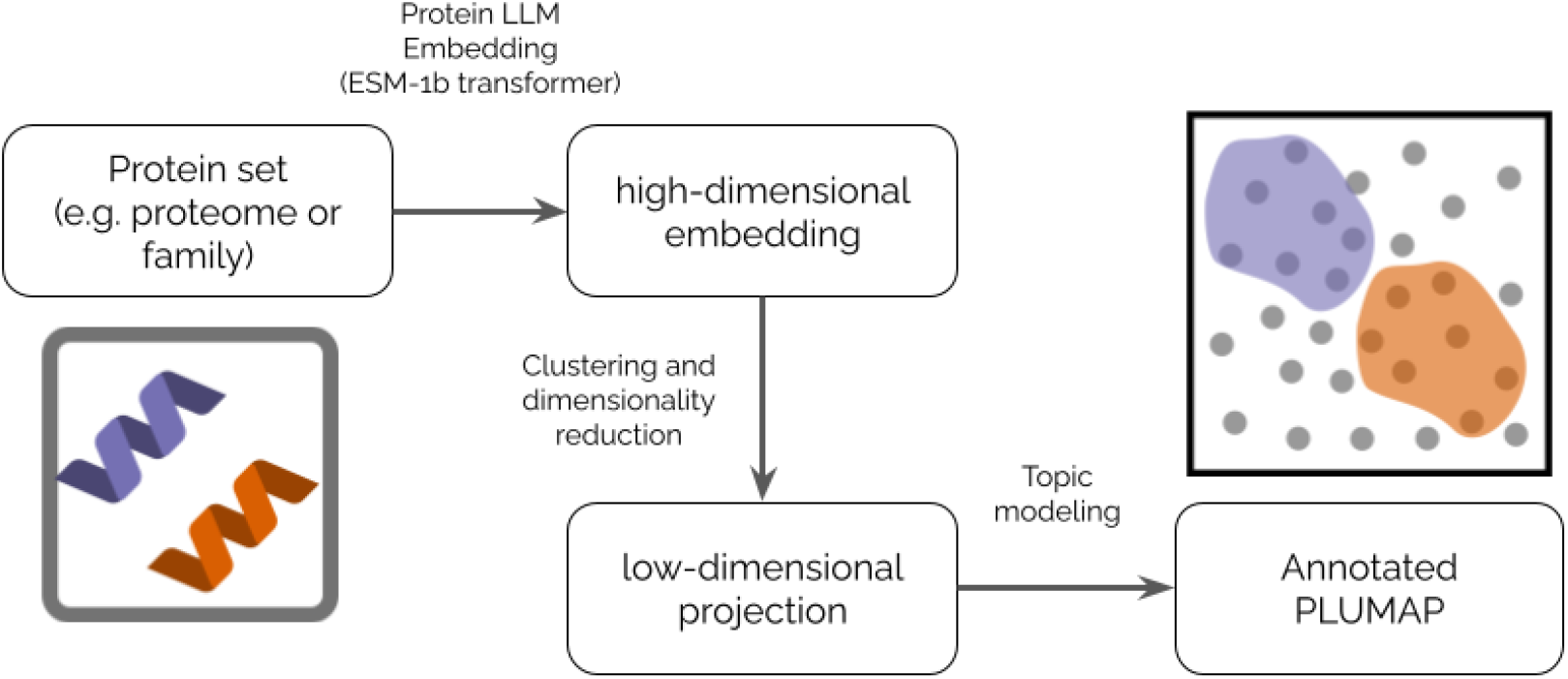
Schematic diagram of steps to generate protein language UMAPs (PLUMAPs) annotated with topic modeling.

**Fig. 2:**
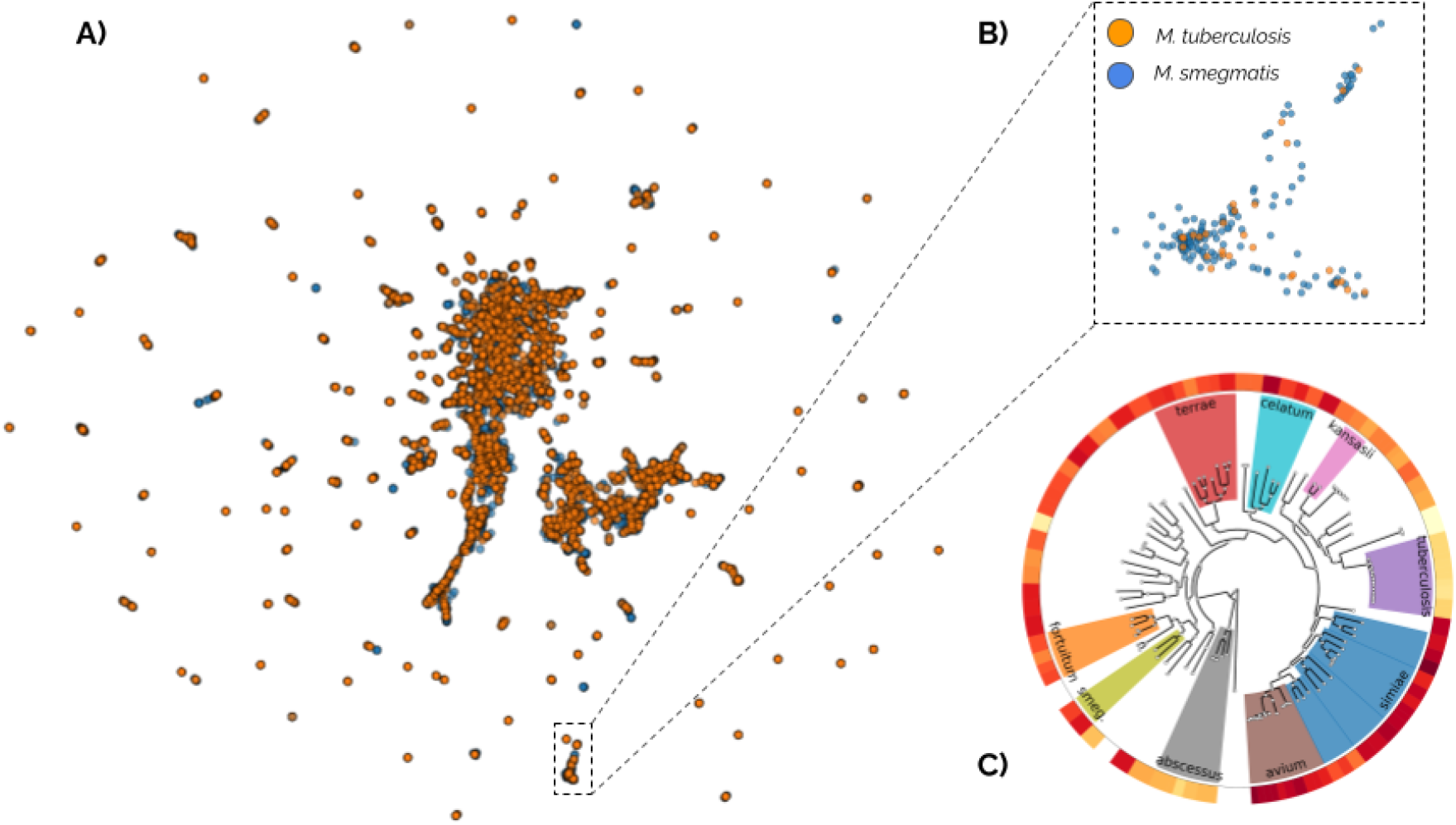
A protein language UMAP (PLUMAP) reveals comparative protein family enrichment between different mycobacterial species. A) 2-dimensional representation of the full proteomes of *M. tuberculosis* (orange) and *M. smegmatis* (blue). Each dot is a single protein. The 2D coordinates embeddings were obtained by dimensionality reduction (via UMAP) of the protein embeddings (vector representations) obtained with the ESM-1b protein language model. B) A cluster of short-chain dehydrogenase/reductase (SDR) family proteins is enriched for *M. smegmatis* proteins. C) Normalized relative abundance of proteins with NAD(P)-binding domains across a diverse set of mycobacterial species.

We also find instances where inspection of a protein cluster hints at the possible function of unannotated enzymes within that cluster. For example, the *Msm* protein MSMEG_5205 (UniProt ID: A0R2R4) of unknown function (UniProt annotation score = 1/5), falls in a cluster enriched for diacylglycerol O-acyltransferases. While the InterPro entry doesn’t contain any informative protein family information, its closest homologue in Mtb, Rv1125, contains an O-acyltransferase WSD1-like domain, supporting the visually generated hypothesis.

### PLUMAPs display protein similarity at multiple scales

Sequence similarity networks (SSNs) have proved to be extremely helpful for exploring homology relationships between proteins ^3–6^. We argue that one advantage of PLUMAPs over SSNs is that they are better at displaying, in one shot, the global structure of protein similarity. More specifically, in addition to conveying that a protein falls *within* a cluster, the distances *between* protein clusters are also informative. In contrast, since the edges connecting proteins in an SSN depend on a threshold stringency value, the relative position of disconnected groups has no meaning; only the observation that a protein falls within a cluster is interpretable. This drawback is addressed by generating multiple SSNs, each one with a different stringency threshold, which is cumbersome. Thus PLUMAPs bring into sharper focus relationships between separate protein clusters, leading to a more efficient visual analysis.

Visually inspecting the layout of protein clusters in the *Mtb/Msm* composite PLUMAP, we find many instances supporting this notion, where protein similarity is displayed at multiple scales (Fig. 3). Two ways in which this is discernible is by the presence of clusters that are near each other in the 2D UMAP, and in clusters that, upon closer inspection, are composed of two or more subclusters. For example, a cluster enriched for short-chain dehydrogenase/reductase (SDR) enzymes is close to two clusters enriched for SAM-methyltransferases and alcohol-dehydrogenases, respectively. The nearby position of the three clusters in the UMAP reflects the fact that all three families have a Rossman fold, which specifies binding to the cofactors NAD(H), NADP(H), and/or SAM. A second example consists of two contiguous clusters, one enriched for acetyl-CoA acyltransferases and thiolases, and the second enriched for enoyl-CoA hydratases and acyl-CoA thioesterases. These two families have HotDog superfamily domains, and are related by their activity on coenzyme-A activated compounds ^14,15^. A third example shows three neighboring clusters enriched for different transcription factor families, the TetR, PadR, LuxR, and merR families. As a fourth example, we note a cluster of proteins with ABC transporter-like, ATP-binding domains, which upon closer inspection, consists of several subclusters, highlighting family substructure within this set of proteins.

**Fig. 3:**
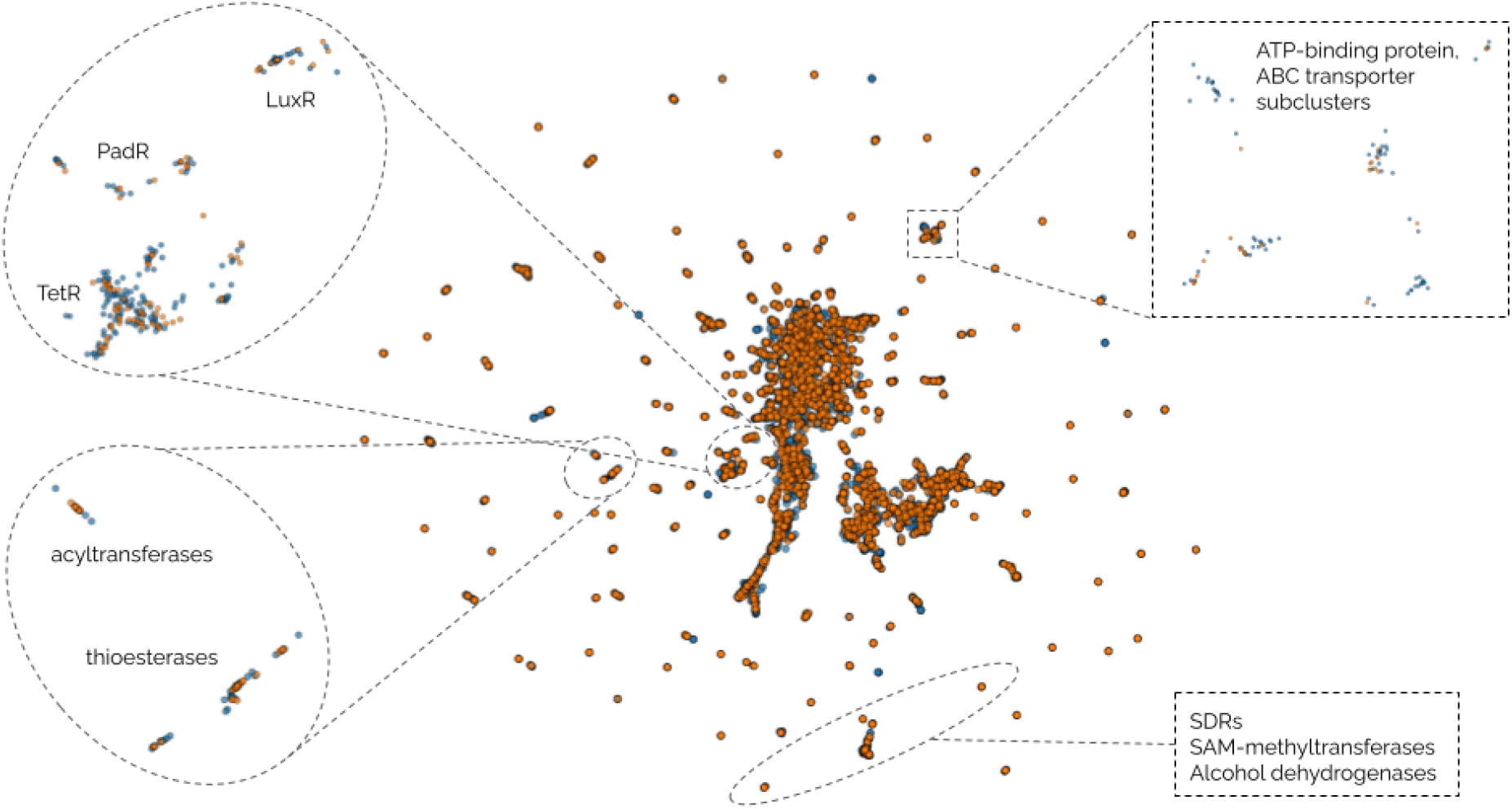
Protein language UMAPs (PLUMAPs) capture protein similarity at different scales. Four regions of the composite *M. tuberculosis*/*M. smegmatis* PLUMAP are highlighted, illustrating how (1) related protein clusters are mapped near each other in the UMAP, and (2) upon closer inspection, some clusters have substructure and are composed of multiple subclusters.

### PLUMAPs group proteins by cellular localization

One of the most striking, bird’s-eye view features of the composite *Mtb/Msm* PLUMAP is the presence of two large, central clusters. Overlaying available protein localization information from UniProt we find that one of the two large clusters is strongly enriched for proteins annotated as localized to the cell wall or membrane (Fig. 4) We find that this grouping by cellular localization is robust across different values of the hyperparameters involved in the PLUMAP pipeline. This illustrates a different way in which PLUMAPs efficiently display protein similarity at different scales, grouping protein sequences with cell wall or membrane-related folds together on average.

**Fig. 4:**
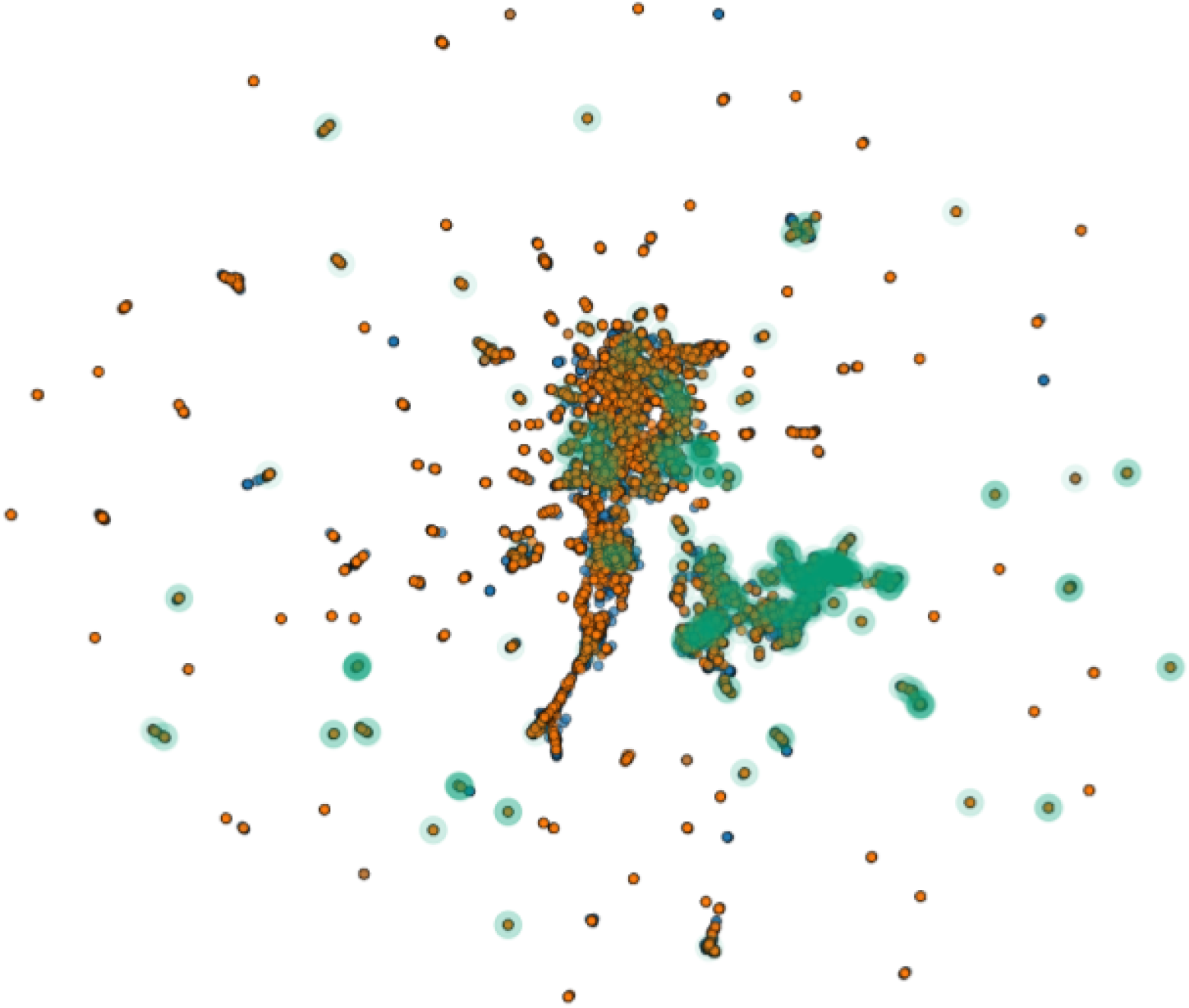
PLUMAPs group proteins by cellular localization. Protein localization information was obtained from UniProt for the Mtb and Msm proteomes. Proteins with the terms “membrane” or “wall” or “lipid” in their localization description are shown in green.

### PLUMAPs as topic modeling: overlaying additional information layers on PLUMAPs

In Natural Language Processing (NLP), topic models aim to detect abstract “topics” in document collections ^13^. Thus, in analogy to topic modeling methods like BERTopic ^10^ we frame PLUMAPs as a protein centered topic modeling technique that leverages protein representations and dimensionality reduction algorithms to create interpretable, and potentially insightful, protein clusters. One way to represent a protein cluster’s “topic” is through enrichment analysis of gene or protein function words or phrases from UniProt annotation tables. We implement this with the Natural Language Toolkit (NLTK) ^16^, a Python library for NLP. Specifically, for all proteins within a cluster, we obtain the frequency distribution of n-grams in the “Protein Name” UniProt entries, which provide an exhaustive list of protein names, information on the activity of the protein, description of the catalytic mechanism of enzymes and functional domains ^1^. An alternative way to explore a cluster’s “topic” is through protein structures, displaying a sample of AlphaFold2 protein structures in a selected cluster. We find that this is pedagogically helpful, since by interactively studying the predicted structures of many proteins in a cluster one can detect common structural patterns, which is more instructive than visualizing a single protein family structure.

## Discussion

Inspired by the practical value of Sequence Similarity Networks (SSNs) ^3–5^, in this work we developed an analogous approach based on protein language models called PLUMAPs. We envision PLUMAPs will be useful in all applications where SSNs have been valuable, in particular for visual analyses of sequence-function relationships.

The design of PLUMAPs is modular, with each component of the pipeline (Fig. 1) replaceable with other equivalent tools. For example, the ESM-1b module could be substituted with a different protein language model, such as ProteinBERT ^9^ or the recent and significantly larger (15-billion parameter) ESM2 model ^8^. Dimensionality reduction could be implemented with other algorithms besides UMAP; and topic modeling could be implemented with other protein language modeling tools, such as Latent Dirichlet Allocation (LDA). Scaling the number of proteomes and entire protein families with pre-computed language model embeddings will speed up usage and analysis.

## Methods

Here, we outline the workflow to create our interactive proteome explorer from transformed vector representations of proteins. To make our tool more accessible to users, we have also created a Colab Notebook which is also referred to in our workflow below.

### Creating a Protein Database: Downloading Proteomes

Proteomes can be downloaded from the UniProtKb database by searching for the organism of interest using the “Proteomes’’ search tool. The results should be mapped to UniProtKb to include sequence and functional information with each protein in the downloaded proteome. Reference proteomes should be used to include more annotated proteins. To use our Colab, proteomes should be downloaded in .tsv format. Alternatively, you can use our preloaded proteomes that have already been transformed.

### Creating a Protein Database: Transforming Proteins to Vector Representations

To create vector representations of proteins in the proteome, we relied on the ESM-1b Transformer available on GitHub ^7^. The ESM Transformer takes as input proteomic data as .FASTA files and outputs 1280-dimensional vector matrices for each protein. We use the final representation layer (No. 33) of the mean representation for each vector for manipulation and visualization.

Our Colab allows users to generate a .FASTA file from multiple proteomes using .tsv files downloaded from UniProtKb which can be input into the ESM transformer. The output directory containing the vector representations can be uploaded to the Colab for computation and visualization.

### Dimensionality reduction with UMAP

To project the matrix representing the proteome into a two-dimensional space, we used the Uniform Manifold Approximation and Projection (UMAP) algorithm, a dimension reduction technique to visualize datasets with high-dimensionality ^14^. We reduce the matrix to three components and use the first two components to project the matrix. To identify potential clusters of proteins that share similarities, we used the K-means clustering algorithm using the sci-kit learn package. The number of clusters was ascertained by optimizing the silhouette coefficient.

### Topic modeling

The clusters were annotated by identifying the most enriched bigram in the protein names within the clusters using the Natural Language Toolkit (Nltk).

